# The C2 domain augments Ras GTPase Activating Protein catalytic activity

**DOI:** 10.1101/2024.08.29.609784

**Authors:** Maxum E. Paul, Di Chen, Kimberly J. Vish, Nathaniel L. Lartey, Elizabeth Hughes, Zachary T. Freeman, Thomas L. Saunders, Amy L. Stiegler, Philip D. King, Titus J. Boggon

**Affiliations:** Department of Molecular Biophysics and Biochemistry, Yale University, New Haven, CT, USA; Department of Microbiology and Immunology; Transgenic Animal Model Core, and; Department of Internal Medicine, University of Michigan Medical School, Ann Arbor, MI, USA; Department of Pharmacology, Yale University, New Haven, CT, USA; Yale Cancer Center, Yale University, New Haven, CT, USA

**Author notes:** These authors contributed equally (Maxum E. Paul, Di Chen).

**Keywords:** p120RasGAP, RASA1, GTP hydrolysis, Ras signaling, GTPase activating protein

## Abstract

Regulation of Ras GTPases by GTPase activating proteins (GAP) is essential for their normal signaling. Nine of the ten GAPs for Ras contain a C2 domain immediately proximal to their canonical GAP domain, and in RasGAP (p120GAP, p120RasGAP; *RASA1*) mutation of this domain is associated with vascular malformations in humans. Here, we show that the C2 domain of RasGAP is required for full catalytic activity towards Ras. Analysis of the RasGAP C2-GAP crystal structure, AlphaFold models, and sequence conservation reveal direct C2 domain interaction with the Ras allosteric lobe. This is achieved by an evolutionarily conserved surface centered around RasGAP residue R707, point mutation of which impairs the catalytic advantage conferred by the C2 domain *in vitro*. In mice, *R707C* mutation phenocopies the vascular and signaling defects resulting from constitutive disruption of the *RASA1* gene. In SynGAP, mutation of the equivalent conserved C2 domain surface impairs catalytic activity. Our results indicate that the C2 domain is required to achieve full catalytic activity of Ras GTPase activating proteins.

## Introduction

The Ras group of small GTPases are essential to a wide variety of signal transduction pathways. These guanine nucleotide-binding proteins cycle between a GDP-bound state, termed the inactive state, and a GTP-bound state which typically binds downstream effector proteins and is thus termed the active state ^1^. Cycling between these states requires either exchange of GDP for GTP, or enzymatic cleavage of the gamma-phosphate of GTP to produce GDP and inorganic phosphate. Both reactions are typically slow without aid of either guanine nucleotide exchange factors (GEF) or GTPase activating proteins (GAP), thus the GEF and GAP proteins are intrinsic to facilitating rapid signaling responses via small GTPase cascades ^1^. The first GAP to be identified was RasGAP (p120GAP, p120RasGAP), encoded by the *RASA1* gene ^2,3^; with nine further GAPs for Ras identified, the group contains ten proteins ^4^.

The role of RasGAP is well established. In mice, loss of functional RasGAP results in dysregulated activation of Ras in endothelial cells leading to impaired developmental, neonatal and pathological angiogenesis and blocked development and maintenance of venous, lymphatic and lymphovenous valves^5-10^. In humans, germline inactivating mutations of the *RASA1* gene cause vascular anomalies, including capillary malformation-arteriovenous malformation (CM-AVM), vein of Galen arteriovenous malformation (VGAM) and lymphatic vessel dysfunction^11-21^. Most identified mutations result in premature stop codons predicted to result in total loss of protein expression from the mutated *RASA1* allele^13-15,19-21^, and combined with mutations in *EPHB4*, the gene for the Ephrin B4 receptor, account for the majority of CM-AVM and VGAM occurrences ^22,23^. However, rare missense germline *RASA1* mutations have also been identified and may provide functional insight into the mechanism of action of RasGAP. Of these mutations, *RASA1 R707C* is found in two families with VGAM^16,17^. This mutation is located in the C2 domain.

C2 domains are commonly thought of as membrane targeting calcium-binding domains, however, these traits are not requisite for this fold and C2 domains often exhibit neither, with the domain playing roles that utilize interactions with partner proteins, or intramolecular interactions, as functional axes ^24^. The RasGAP C2 domain is located immediately N-terminal to its GAP domain and a similarly located C2 domain is found immediately N-terminal to the GAP domain in nine of the ten GAPs for Ras small GTPases, except for neurofibromin (**Fig. 1A**)^4,25,26^. The conserved nature of this domain across the GAPs for Ras is not well understood, but in RasGAP, the GAP domain is insufficient for full catalytic activity^27^ and in SynGAP, the presence of the C2 domain augments catalytic activity against Ras and the related GTPase, Rap^28^. In addition, in other dual-specificity RasGAPs (RASA2, RASA3, RASAL1), the presence of the C2 domain confers a catalytic advantage compared to the GAP domain^29,30^. In RasGAP, other domains have been proposed as activity modulators^27,31-34^ but potentially not a direct impact on activity^35^. Together these datapoints suggest that the C2 domain of RasGAP may have a role in catalytic activity towards Ras.

**Figure 1.**
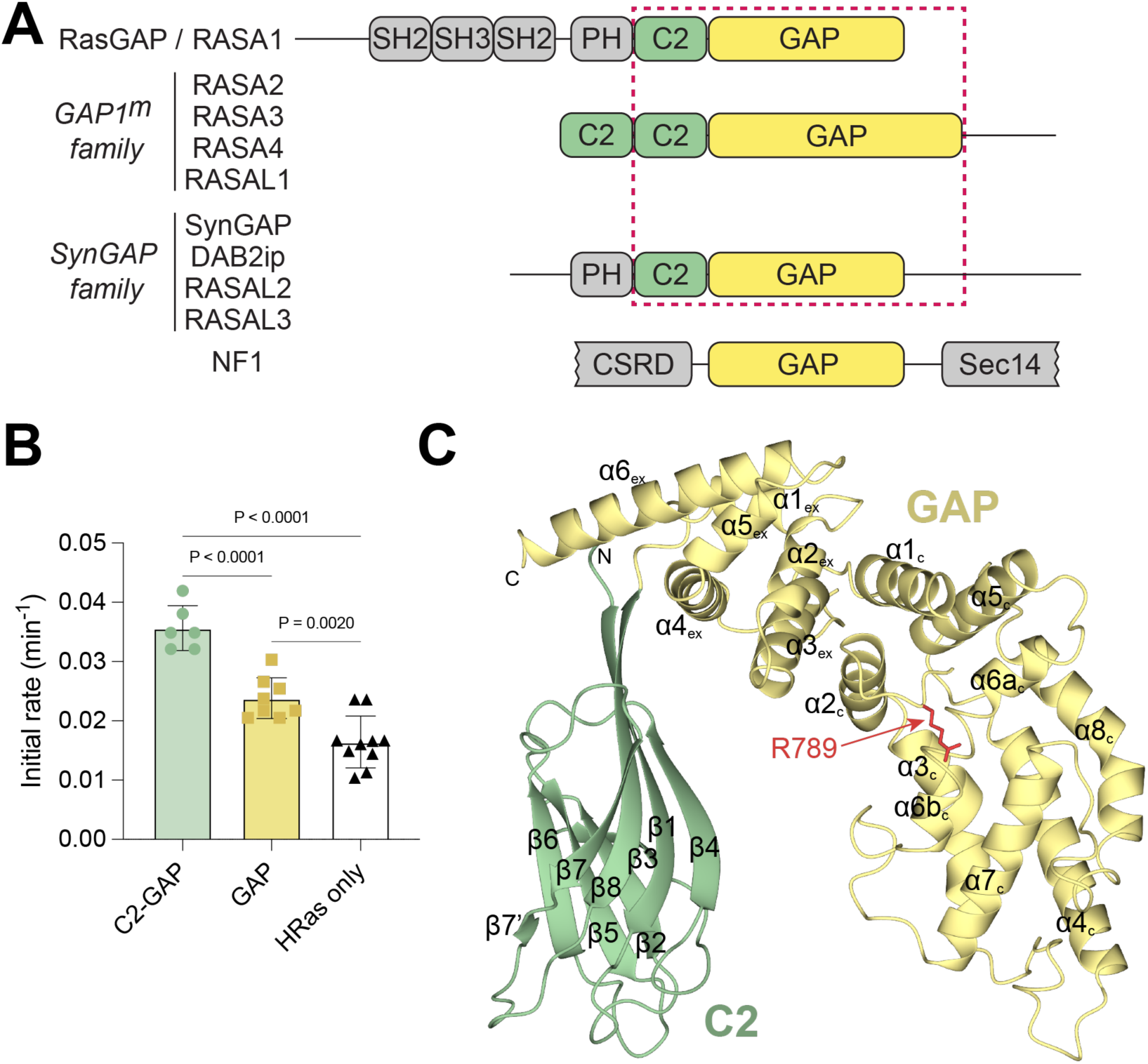
RasGAP C2 domain location and structure. **A)** Schematic architecture of GTPase Activating Proteins for Ras. RasGAP GAP1^m^ family: RASA2, RASA3, RASA4, RASAL1. SynGAP family: SynGAP, Dab2ip, RASAL2, RASAL3. SH2, Src homology 2; SH3, Src homology 3; PH, pleckstrin homology; C2, protein kinase C domain 2; CSRD, cysteine-serine-rich; Sec14, *S. cerevisiae* phosphatidylinositol transfer protein homology. **B)** Single turnover GAP assay. Initial rates for Phosphate Sensor reporting on phosphate release from GTP loaded Ras. RasGAP constructs: C2-GAP, GAP, and HRas control. Bars indicate mean ± SD with replicates (n=6, 8, and 10) shown. P values for relevant comparisons are shown. Statistical significance determined via ordinary one-way ANOVA (*F* = 44.55, 23 degrees of freedom) with Tukey’s multiple comparisons test. GAP activity is increased by the addition of the C2 domain. Assay data shown in Table S1. **C)** Crystal structure of the C2-GAP region of RasGAP. The GAP domain is colored yellow, the C2 domain is colored green. GAP and C2 domain domain secondary structures assigned as previously^36,37^. The arginine finger R789 of the GAP domain is indicated in red. Crystallographic statistics are shown in Table S2. PDB accession code: 9BZ4.

In this study, we assess the role of the C2 domain of RasGAP and reveal that it is required for full catalytic activity towards Ras. We determine the 2.45 Å crystal structure of the C2-GAP domain region of RasGAP and through analysis of conformational states in the asymmetric unit, comparison with previous structures of RasGAP with Ras, AlphaFold models of complexes, and sequence conservation, we reveal that the C2 domain interacts with the Ras allosteric lobe. We find this interaction to be mediated by an extensive conserved surface centered around RasGAP residue R707. Using *in vitro* enzymatic assays, we demonstrate that constructs including the C2 domain are more active towards Ras than the GAP domain alone, but that mutation of R707 impairs the catalytic advantage. To assess the importance of the C2 domain in an animal model we generated the *R707C* mutation in mice. We find that this phenocopies the vascular and signaling defects resulting from constitutive disruption of the *RASA1* gene. Together these results demonstrate that the C2 domain is essential for full RasGAP activity. Strikingly, a C2 domain is present immediately preceding the GAP domain in nine of the ten GAPs for Ras. We compared the residues conserved in RasGAP with these other GAPs for Ras. This revealed high conservation across the nine proteins and evolutionary conservation within each gene, with functional importance confirmed by mutation of the conserved surface in SynGAP C2 domain, which similarly impairs catalytic activity. Overall, our results indicate that the C2 domain is required to achieve full catalytic activity of Ras GTPase activating proteins.

## Results

### The RasGAP C2 domain augments GAP activity

The RasGAP C2 domain is located immediately N-terminal to its GAP domain. To address whether the presence of this domain impacts catalytic activity toward Ras, we conducted *in vitro* multiple turnover GAP assays for RasGAP. These assays revealed that the GAP domain alone is significantly less active than a construct containing the C2 and GAP domains (**Fig. 1B, Table S1**). This prompted us to crystallize the C2-GAP region of RasGAP to determine its structure.

### The structure of RasGAP C2-GAP

We determined a 2.45 Å crystal structure of RasGAP C2-GAP (residues 588-1044) (**Fig. 1C, Table S2**) and observe that the GAP domain is conformationally similar to the structure determined previously for the isolated GAP domain^36^ with root-mean-square deviations ranging between 0.6 Å and 1.4 Å over 324 residues. The type II C2 domain^37^ is most similar to the previously determined structure of the C2 domain of Protein kinase Cη (Dali server^38^) and does not contain canonical calcium binding residues (**Fig. S1**). The crystal structure contains four copies of RasGAP C2-GAP in the asymmetric unit. When superposed onto the catalytic core GAP domain (residues 761-977) determined previously^36^, we found that there is conformational heterogeneity in the location of the C2 domain (residues 590-713) with equivalent carbon alpha atoms located up to 12 Å distal from one another across the four copies (**Fig. 2A, Fig. S2**). As the crystal structure of the RasGAP GAP domain in complex with Ras was previously determined^39^, we superposed the C2-GAP structure and found that the C2 domain may approach as close as 7 Å from Ras (**Fig. 2B**). This intriguing result prompted us to conduct two-chain predictions of RasGAP C2-GAP and Ras using AlphaFold^40,41^. In these predictions we found a further rotation in the C2 domain that suggests direct contact between the C2 domain and the Ras allosteric lobe (**Fig. S3**). Together, these structural analyses and molecular models suggest that the RasGAP C2 domain may directly contact Ras.

**Figure 2.**
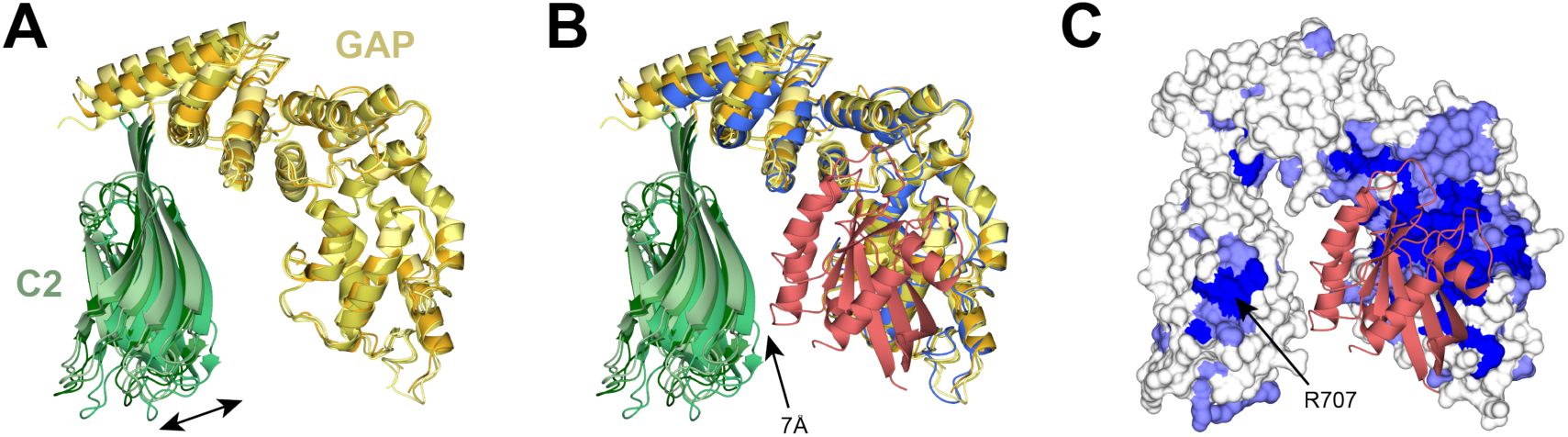
RasGAP C2-GAP structure. **A)** Superposition of the four copies of RasGAP C2-GAP found in the asymmetric unit onto the GAPc sub-domain (residues 761-977^36^), illustrating conformational movements of the C2 domain. **B)** Superposition of the co-crystal structure of RasGAP (blue) with Ras (red) (1WQ1 ^39^) on the core GAP subdomain. Four copies of the C2-GAP crystal structure are colored as in (a). The closest distance between C2 and Ras is indicated. **C)** Evolutionary conservation of RasGAP C2-GAP surface residues. Blue indicates complete conservation, white indicates low conservation. Conservation mapping conducted using the Consurf server^58^. Ras from the superposed complex of Ras with the RasGAP GAP domain is shown in red (1WQ1^39^).

We next asked if this unexpected predicted interaction between the C2 domain and Ras is evolutionarily conserved. We aligned 209 sequences of RasGAPs from humans to sponges (**Fig. S5**), mapped conservation onto the structure (**Fig. S6**) and superposed the Ras-bound GAP domain^39^ (**Fig. 2C**). As expected, we found complete conservation of the Ras-binding region in the GAP domain (**Fig. 2C**). However, unexpectedly, in the C2 domain we found the surface predicted to be in closest proximity to Ras is also completely conserved from humans to sponges (**Fig. 2C**). This contiguous patch comprises residues S705 and R707 of strand β8, D687 and W689 of strand β7, and H604 of strand β1 (**Fig. S5**). We posited that the high conservation through evolution indicates a functionally important role for this C2 domain surface.

### Mutation of the RasGAP C2 domain impairs catalytic advantage

Residue R707 is both central in the conserved C2 domain surface and predicted by AlphaFold to directly interact with Ras (**Fig. S3**). Mutation of this residue (R707C) has also been reported in patients with VGAM^16,17^. Therefore, we asked if mutation of R707 affects RasGAP catalytic activity. We evaluated the impact of R707 mutation by assessing the Michaelis-Menten kinetics of RasGAP towards Ras with a single turnover phosphate release assay using the Phosphate Sensor system to detect inorganic phosphate. Compared to the GAP domain alone, C2-GAP exhibited significantly increased catalytic efficiency, which is primarily driven by an increased k_cat_ (**Fig. 3A-C, Table S3**). Moreover, mutations R707S or R707D in the C2-GAP construct abrogated the gain in catalytic efficiency achieved by the presence of the C2 domain. This finding is also observed for longer constructs of RasGAP harboring R707S, R707C, and W689F mutations (**Figure S6, Table S4**).

**Figure 3.**
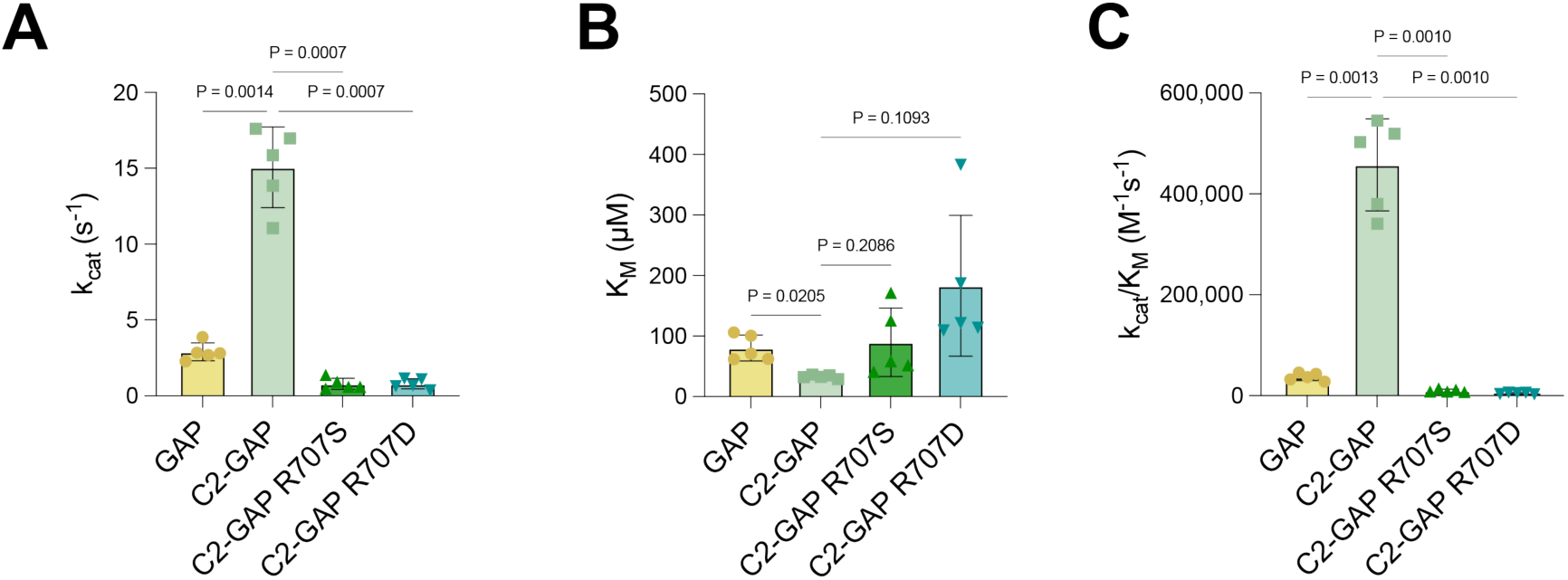
Activity profile of RasGAP. **A-C)** Michaelis-Menten parameters from single turnover phosphate release assays. k_cat_, K_M_, and catalytic efficiency (k_cat_/K_M_) of RasGAP constructs are shown. Bars indicate mean ± SD (n=5). *P* values for relevant comparisons are shown. Statistical significance was determined with the Brown-Forsythe ANOVA test (A: *F* = 123.2, 7.686 degrees of freedom; B: *F* = 4.585, 9.113 degrees of freedom; C: *F* = 116.1, 7.065 degrees of freedom) with Dunnett’s T3 multiple comparisons test.

### Impaired vascular development in RasGAP R698C mice

To assess the functional significance of the predicted RasGAP C2 domain interaction with Ras *in vivo*, we used CRISPR/Cas9 targeting to generate a *Rasa1 R698C* allele in mice, which is equivalent to human *RASA1 R707C*. Heterozygote *Rasa1 R698C* mice were viable and fertile. However, no *Rasa1 R698C* homozygous pups were identified in litters from crosses of heterozygous parents, indicating that the *Rasa1 R698C* allele is embryonic lethal in heterozygous form **(Fig. 4A)**. To determine the time of embryonic lethality, we set up timed matings. At E9.5, *Rasa1 R698C* homozygous embryos were externally indistinguishable from wild type and *Rasa1 R698C* heterozygote littermates and were recovered in ratios expected for normal Mendelian inheritance **(Fig. 4B)**. However, by E10.5, the majority of homozygote embryos were abnormally small (6/9 embryos) and exhibited grossly distended pericardial sacs or evidence of cutaneous hemorrhage (2/9 embryos), consistent with abnormal cardiovascular development. At E11.5-12.5, of only two homozygotes recovered (2/19 total embryos), both showed distended pericardial sacs, and size differences compared to controls were accentuated. No homozygous *Rasa1 R698C* homozygous embryos were recovered after E12.5. Levels of expression of RasGAP protein at E9.5 were not affected by the R698C mutation, indicating that the developmental abnormalities were not a result of reduced abundance of RASA1 **(Fig. 4C)**. As a control, we generated mice that carry a *Rasa1 T603A* mutation that is also located in the C2 domain. T603A is equivalent to human *RASA1 T612A* that has been reported in a family with AVM^42^. The mutation resides in the loops of the C2 domain distant from the interface containing R707. In litters of crosses of *Rasa1 T603A* heterozygous parents, *Rasa1 T603A* homozygous pups were present in the expected Mendelian ratio (**Fig. S4**). Thus, the *Rasa1 T603A* mutation is not embryonic lethal in homozygous form and not all mutations of the RasGAP C2 domain cause equivalent phenotypes *in vivo*.

**Figure 4.**
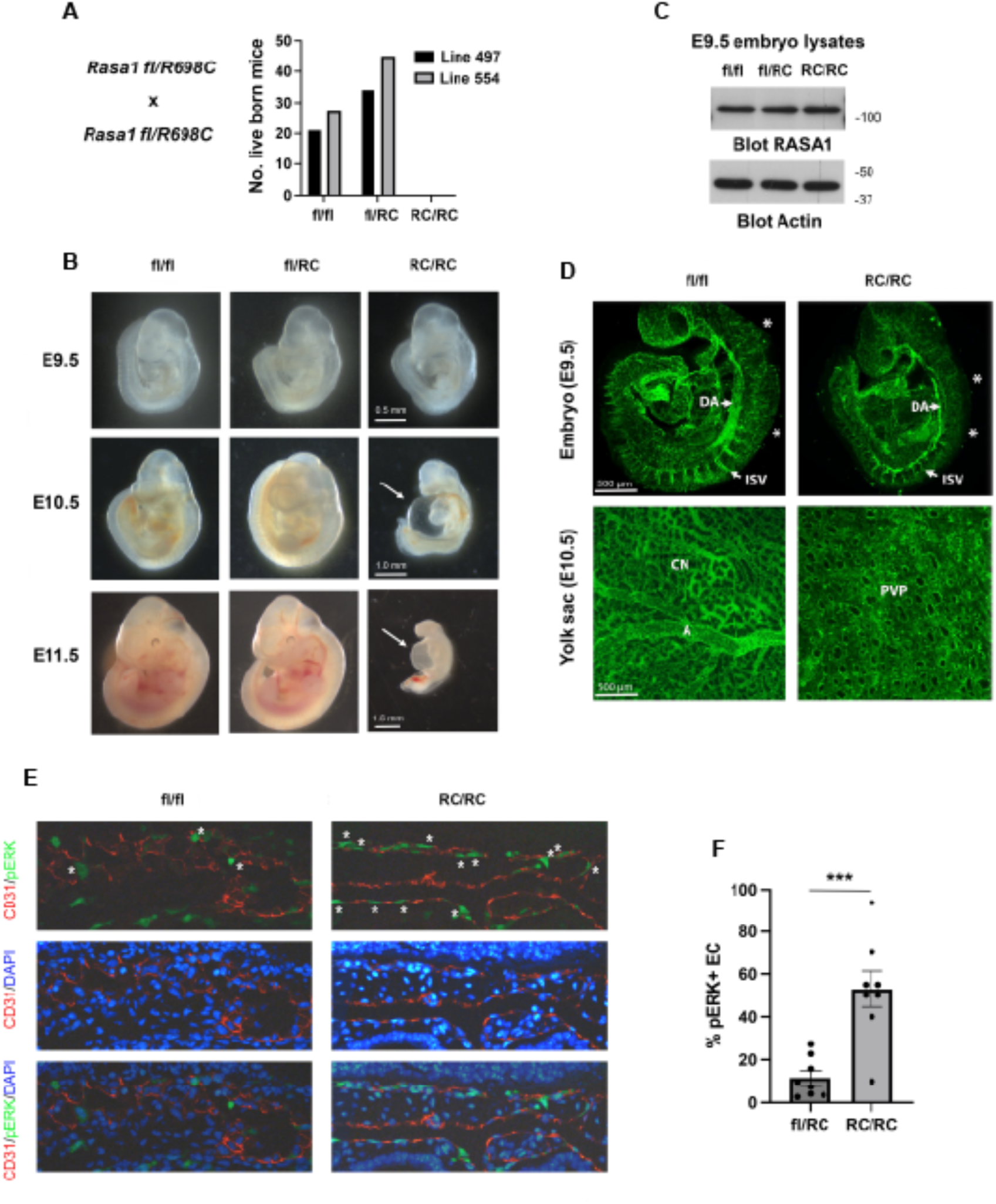
Embryonic lethality and vascular defects in homozygous *Rasa1 R698C* mice. *Rasa1 fl/R698C* heterozygous mice derived from two independent founders (lines 497 and 554) were crossed. **A)** Numbers of live-born pups of the indicated genotypes are shown for each line. *P*<0.0001, Chi-squared test. **B)** Embryos at different stages of development were harvested from timed matings. Note smaller sizes of *Rasa1 R698C* homozygote embryos with distended pericardial sacs (arrows) at E10.5 and E11.5. **C)** Amounts of RASA1 and actin in E9.5 whole embryo lysates were determined by Western blotting. **D)** E9.5 embryos and E10.5 yolk sacs were stained with CD31 antibodies to highlight the vasculature. Note an irregular dorsal aorta (DA) and intersomitic vessels (ISV), and incomplete developmental angiogenesis toward the midline (asterisks) in the *Rasa1 R698C* homozygote embryo and a primitive vascular plexus (PVP) in the yolk sac that is distinct from a remodeled vasculature comprising of a capillary network (CN) that drains to arteries (A) in the control yolk sac. **E)** Sections of E9.5 embryos were stained with antibodies against CD31 and phospho-active ERK MAPK (pERK). Images show part of the dorsal aorta. Some pERK+ CD31 are indicated with asterisks. **F)** The percentage of aortic pERK+ EC in non-overlapping microscopic fields (as in e) was determined. Shown is the mean +/- 1SEM of the percentage of pERK+ EC. *** *P*<0.001, two-sided Mann Whitney test.

To examine vascular development in *Rasa1 R698C* homozygotes, we stained E9.5 embryos and E10.5 yolk sacs with CD31 antibodies **(Fig. 4D)**. At E9.5, *Rasa1 R698C* homozygote embryos exhibited an irregular dorsal aorta and intersomitic vessels and impaired angiogenesis toward the midline compared to littermate controls. At E10.5, the yolk sac contained a primitive vascular plexus that was not remodeled into a hierarchical vascular network comprising of capillaries and arteries that deliver blood to the developing embryo, as in controls. These abnormalities of vascular development have been identified previously in constitutive RasGAP-deficient embryos and embryos that express a catalytically-inactive form of RasGAP with a point mutation in the arginine finger of the GAP domain (R780Q), also at E9.5 and 10.5^6,8^. Therefore, the RasGAP R698C mutation phenocopies loss of RasGAP and RasGAP R780Q mutations in mice, consistent with an essential role for RasGAP C2 domain interaction with Ras for normal Ras regulation *in vivo*. To confirm that Ras-mitogen-activated protein kinase (MAPK) signaling was dysregulated in *Rasa1 R698C* homozygous embryos, sections of E9.5 embryos were stained with antibodies against phospho-active ERK MAPK and CD31. Compared to control embryos, *Rasa1 R698C* homozygous embryos showed significantly higher levels of phospho-MAPK in EC of the dorsal aorta **(Fig. 4E,F)**.

### The C2 domain augments catalytic activity across GAP proteins for Ras

We used AlphaFold to predict the structure of the C2 and GAP domains of all other C2 domain-containing RasGAPs. In each case, the GAP-proximal C2 and GAP domains were oriented similarly to those in RasGAP and SynGAP C2-GAP^28^ (**Fig. S8**). Sequence conservation for each protein over evolution was mapped onto the surface of each AlphaFold model and revealed conserved surfaces in each of the GAP-proximal C2 domains. These surfaces encompassed a region similar to the conserved surface observed in the RasGAP C2 domain (**Fig. 5A, Fig. S9**). We next asked if the amino acid residues comprising this conserved surface resemble one another, and on analysis found them to be almost identical to those in RasGAP, with R707 completely conserved across the family except in RASA4 (**Fig. 5B, Fig. S10**). To assess the importance of this highly conserved surface we introduced an R401 mutation into SynGAP, equivalent to R707 in RasGAP, and found that it completely abrogated catalytic activity towards its preferred substrate Rap1 (**Fig. 5C, Table S5**). It is unlikely that this level of surface conservation across nine members of a protein family and throughout evolution for each family member is necessary for an adaptor function for the C2 domain. Instead, we propose that the conservation reflects an important role of this surface in the regulation of Ras across all nine C2 domain-containing family members. Therefore, we conclude that catalytic activity for the Ras GTPase Activating Protein family is augmented by a surface patch conserved in each of the GAP domain-proximal C2 domains that directly interacts with the Ras allosteric lobe.

**Figure 5.**
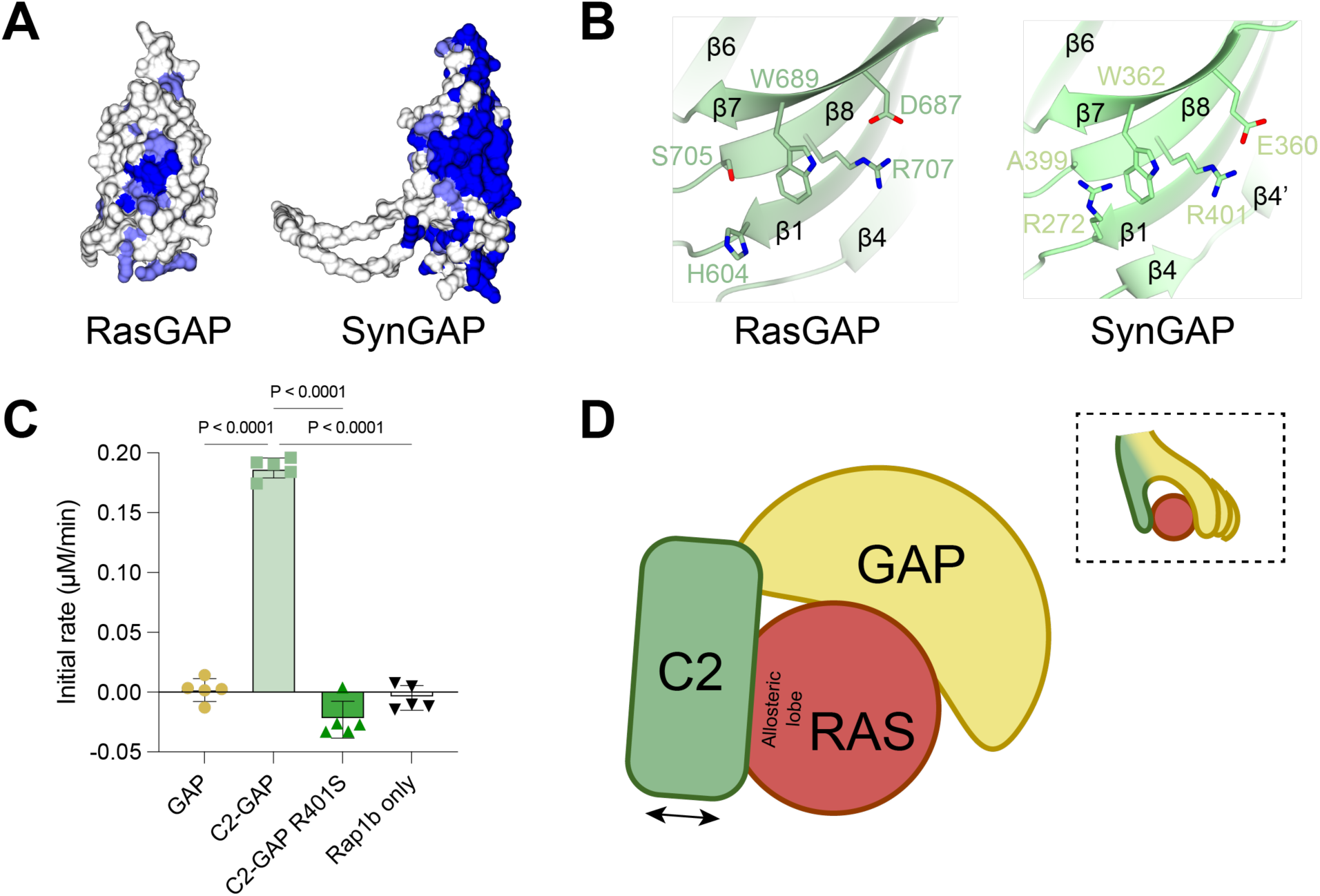
Conservation of the role of the C2 domain. **A)** Conservation of the C2 domain of RASA1 and the AlphaFold-predicted GAP-proximal C2 domain of SynGAP; AF-Q96PV0-F1. Shown is the front face of the C2 beta sandwich, which comprises strands β7, β8, β1, and β4. High conservation is colored blue, low conservation is white. **B)** Closeup of evolutionarily conserved C2 domain residues in RASA1 (left) and SynGAP (right). Secondary structure features are indicated. Both proteins share a conserved arginine (β8) and phenylalanine (β7) on the C2 face, as well as conserved acidic residues on strand β7, positive residues on strand β1, and small uncharged residues on β8. **C)** SynGAP multiple turnover activity assay in the presence of 5µM Rap1b. Bars indicate mean ± SD with replicates (n=5) shown. *P* values for relevant comparisons are shown. Statistical significance was determined via ordinary one-way ANOVA (*F* = 385.7, 19 degrees of freedom) with Dunnett’s multiple comparisons test. Rap1b GTPase activity is stimulated in the presence of SynGAP C2-GAP, but is not detectable with SynGAP GAP, SynGAP C2-GAP R401S, or without SynGAP (intrinsic rate). **D)** Cartoon depicting proposed mechanism of C2 domain regulation of Ras activity, inset illustrating the C2 domain acting as an opposable thumb.

## Conclusions

Our findings show an important role for the GAP-proximal C2 domain in RasGAP in Ras cycling. In structural analyses, we find the C2 domain of RasGAP is positioned to contact the Ras allosteric lobe, and in enzymatic assays, its presence augments catalytic activity. Sequence analyses of RasGAP C2 domains throughout evolution reveal a highly conserved patch centered around residue R707 that is implicated in the augmentation of Ras cycling, which we confirmed by mutagenesis. In a mouse model, we find that constitutive expression of RasGAP R698C (equivalent to R707C) phenocopies mice with constitutive loss of RasGAP or constitutive expression of catalytically inactive RasGAP, with embryonic lethality associated with failed developmental blood vascular angiogenesis and increased MAPK activation in endothelial cells at E9.5-E10.5. Remarkably, each of the C2 domains in the SynGAP and GAP1^m^ subfamilies demonstrate surface conservation in a region similar to that found in RasGAP, with the residues of this conserved surface almost identical across all nine proteins. Similar to RasGAP, an intact C2 domain surface is required for maximal GAP activity of SynGAP. Our study, therefore, indicates that the conserved C2-GAP architecture may be required for normal GAP regulation of Ras signaling across nine of the ten GAPs for Ras.

Our finding that the C2 domain is important for the catalytic activity of RasGAPs predicts that C2 domain mutations that disrupt interaction with Ras should be associated with at least some diseases that are driven by dysregulated Ras activation. Consistent with this, the RasGAP R707C mutation has been reported in VGAM^17,23^, and the SynGAP W362R mutation (equivalent to RasGAP W689) causes non-syndromic intellectual disability and fails to control MAPK activation^43,44^. Likewise, less well studied but clinically observed mutations also exist in cancer, where mutation of the conserved arginine occurs in seven of the nine C2 domain-containing GAPs for Ras (**Table S6**). We posit that disruption of the C2 domain interaction with Ras may prove to be clinically significant for the majority if not all of these GAP proteins for Ras small GTPases.

We provide new insights into the role of the C2 domain, but this study does not reveal the molecular mechanism by which the C2 domain augments Ras hydrolysis. One possibility is that inorganic phosphate release is enhanced. We find that Ras K_M_ is only moderately impacted by the presence of the C2 domain, but that k_cat_ is increased. This correlates with previous studies demonstrating that cleavage of the bond between the β and γ phosphates of GTP occurs almost concurrently with GAP binding and that the rate limiting step for GAP-mediated hydrolysis is release of inorganic phosphate^45-47^. We propose that impingement of the C2 domain upon the Ras allosteric lobe increases k_cat_ by accelerating release of inorganic phosphate.

The role of the C2 domains across the GAPs for Ras has not been extensively studied. By assessing the prototypical GAP, RasGAP, we reveal new insights important for understanding how Ras signaling is controlled. A conserved interaction of the C2 domain with the allosteric lobe of Ras, akin to an opposable thumb (**Fig 5D**), provides new mechanism for how Ras cycling is optimized and new avenues for understanding developmental biology, disease mechanisms and potentially for identifying therapeutic approaches.

## Supplemental Information

Figures S1 to S10 and Tables S1 to S6

## Methods

### Protein expression and purification

Protein constructs were sub-cloned to generate recombinant expression vectors encompassing the following residues in human RasGAP (UniProt ID: P20936), HRas (UniProt ID: P01112), SynGAP (UniProt ID: Q96PV0) and Rap1b (UniProt ID: P61224). RasGAP ΔN construct encompasses residues 174-1047, RasGAP C2-GAP residues 582-1047, RasGAP GAP residues 714-1047, HRas residues 1-167, SynGAP C2-GAP residues 244-740, SynGAP GAP residues 408-740 and Rap1b encompasses residues 1-167. RasGAP ΔN, RasGAP C2-GAP, RasGAP GAP, HRas were cloned into a modified pET-32 (Novagen) bacterial expression vector coding for a hexa-histidine (His_6_) tag followed by a TEV (tobacco etch virus) protease cleavage site using BamHI and XhoI restriction sites. Rap1b cDNA was subcloned into pCDFDuet-1 (EMD Millipore). SynGAP expression constructs were codon-optimized for *E. coli* expression, and synthesis and subcloning into pET-28a(+)-TEV (GenScript). Site-directed mutagenesis was performed using the QuikChange protocol (Agilent) and all mutants except C2-GAP R707D were purified using the same protocol as their corresponding wild type constructs. RasGAP ΔN contains four cysteine to serine mutations (C261S, C236S, C372S, C402S) to prevent multimerization as described in ^48^.

All plasmids were transformed into Rosetta (DE3) cells, which were grown in Luria Broth at 37°C to A_600_ of 0.5-0.7. Expression was then induced with 0.2 mM isopropyl β-D-thiogalactopyranoside (IPTG) at 18°C overnight. Cells were pelleted at 2000 rcf for 30 minutes at 4°C, resuspended in 10 mL lysis buffer (500 mM NaCl, 50 mM HEPES pH 8), and lysed by addition of 50 µg/mL lysozyme, three freeze-thaw cycles, and sonication. Following centrifugation at 48,000 rcf for 1 hour at 4°C the supernatant was applied to 1 mL Ni-NTA agarose beads (ThermoFisher Scientific) and rocked for 1 hour at 4°C. Beads were washed with lysis buffer containing imidazole at concentrations of 20 mM, 40 mM, 100 mM, 250 mM, and 500 mM, fractions containing 40-500 mM imidazole were pooled and His_6_ tagged TEV protease added. Following dialysis overnight at 4°C against 1 L lysis buffer in dialysis tubing with a 10 kDa cutoff. The mixture was again incubated for 1 hour at 4°C on 1 mL Ni-NTA agarose beads to capture uncleaved protein. Flow through and washes containing 20 mM and 40 mM imidazole were pooled and concentrated in an Ultra Centrifugal Filter (Amicon) with a 30 kDa cutoff to a volume of approximately 2 mL, diluted to 25 mL in 20 mM Tris pH 8.5, and then applied to a MonoQ column (GE Healthcare) to perform anion exchange chromatography using a continuous NaCl gradient from 0-40% of 1 M NaCl in 20 mM Tris pH 8.5. The eluted peak was concentrated and subjected to size exclusion chromatography on a Superdex 75 Increase 10/300 GL (GE Healthcare). Samples eluted as monodisperse peaks, and relevant fractions were combined and concentrated. RasGAP samples were diluted in 50% glycerol and stored at -20°C until use. HRas and Rap1b were frozen in liquid nitrogen and stored at -80°C until use. Sample identity and integrity were monitored using SDS-PAGE throughout purifications.

SynGAP GAP and SynGAP C2-GAP were prepared as described above but were not cleaved with TEV protease and did not undergo anion exchange chromatography. Instead, samples from imidazole elutions were concentrated and subjected to size exclusion chromatography directly. Samples eluted as monodisperse peaks, and relevant fractions were combined, concentrated, frozen in liquid nitrogen, and stored at -80°C until use. Sample identity and integrity were monitored using SDS-PAGE throughout purifications.

RasGAP C2-GAP R707D was cloned into a pGEX-6P-1 vector (Cytiva) coding for glutathione S-transferase followed by a PreScission protease cleavage site using BamHI and XhoI restriction sites. Cells were transformed, induced, pelleted, and lysed as above using an alternate lysis buffer (150 mM NaCl, 20 mM Tris pH 8). Following pelleting at 48,000 rcf for 1 hour at 4°C, the supernatant was applied to 0.5 mL Glutathione Sepharose 4B beads (Cytiva) and incubated for 2 hours at 4°C. Beads were washed with 50 mL buffer to remove unbound proteins. Protein was then cleaved on-bead via addition of GST tagged PreScission protease at 4°C overnight. The following day, cleaved protein was washed off in 6 mL buffer, diluted to 28 mL in 20 mM Tris pH 8.5, and then applied to a MonoQ column (GE Healthcare) to perform anion exchange chromatography using a continuous NaCl gradient from 0-40% of 1 M NaCl in 20 mM Tris pH 8.5. The eluted peak was concentrated and subjected to size exclusion chromatography on a Superdex 75 Increase 10/300 GL (GE Healthcare). Samples eluted as monodisperse peaks and were concentrated, diluted in 50% glycerol, and stored at -20°C until use. Sample identity and integrity were monitored using SDS-PAGE throughout purifications.

### Crystallography

Purified RasGAP C2-GAP was extensively screened at 20 mg/mL protein concentration by vapor diffusion methodology in sitting drops at room temperature using an NT8 liquid handling robot (Formulatrix). Using the PEG/Ion HT screen (Hampton Research) initial thin needles were observed in a condition containing 0.2 M ammonium phosphate dibasic and 20% w/v PEG 3350 with a 1:1 v/v protein:reservoir solution ratio in a final drop volume of 0.5 µL. Optimized crystals were obtained using streak seeding in hanging drop vapor diffusion methodology after two days of equilibration at room temperature where drops contained a 1:1 v/v protein:reservoir solution ratio of protein at 10 mg/mL protein and a reservoir solution containing 0.2 M ammonium phosphate dibasic, 0.1 M MOPS pH 7.4, and 26% w/v PEG 3350 in a final drop volume of 2 µL. Crystals were cryopreserved in 0.2 M ammonium phosphate dibasic, 0.1 M MOPS pH 7.4, 26% w/v PEG 3350, 28% ethylene glycol, and flash frozen in liquid nitrogen. Data were collected at beamline 24-ID-C at the Northeastern Collaborative Access Team (NE-CAT) facility at the Advanced Photon Source at Argonne National Laboratory, IL. A single crystal was subjected to three 180° data collection sweeps before succumbing to radiation damage. Data from all three runs were processed and merged using XDS (v20220110)^49^ and CCP4 (v7.0.048)^50^. Matthew’s probability calculation predicted four copies of RasGAP C2-GAP in the asymmetric unit. The merged dataset was processed in space group *P* 1 with unit cell dimensions a=70.9 Å, b=94.0 Å, c=96.7 Å, α=95.8°, β=111.1°, γ=108.5° at a resolution range of 2.45-87.52 Å. Molecular replacement was performed using Phaser (v2.8.3)^51^ in the Phenix (v1.20.1) software package^52^. Phaser search ensembles were generated from the AlphaFold Protein Structure Database^40^ prediction for Ras GTPase-Activating Protein 1 (accession code: AF-P20936-F1) residues 590-713 for the C2 domain, residues 714-763 and 982-1042 for the GAPex sub-domain, and residues 764-981 for the GAPc sub-domain, with GAPex and GAPc as defined by ^36^. Phaser yielded a final overall TFZ score of 30.1 and placed four copies of RasGAP C2-GAP each consisting of three search ensembles with good placement that resulted in unambiguous connectivity between the chains. Phenix (v1.20.1) Autobuild^53^ was performed and cycling between manual building in Coot (v0.9.8.2)^54^ and Phenix (v1.20.1) Refine^55^ was used for ten successive rounds of refinement to obtain the final crystal structure. Assessment of structural quality was performed on the MolProbity server ^56^. The completed structure has been deposited to the Protein Data Bank under accession code 9BZ4 and diffraction images to SB Grid Data Bank under accession code 1107.

### Molecular models using AlphaFold

Monomeric AlphaFold structure predictions were obtained from the AlphaFold Protein Structure Database^40^. Accession codes used for analysis were as follows: for RasGAP, AF-P20936-F1; RASA2, AF-Q15283-F1; RASA3, AF-Q14644-F1; RASA4, AF-O43374-F1; RASAL1, AF-O95294-F1; RASAL2, AF-Q9UJF2-F1; RASAL3, AF-Q86YV0-F1; SynGAP, AF-Q96PV0-F1; DAB2ip, AF-Q5VWQ8-F1. AlphaFold multimer predictions were generated using ColabFold v1.5.5 (AlphaFold2 using MMseqs2) in either Google Colab^41^ or ChimeraX v1.5^57^ using default parameters and primary sequences of HRas (residues 1-167) with either RasGAP C2-GAP (residues 582-1047), PH-C2-GAP (residues 470-1047), or ΔN (residues174-1047).

### Structural alignments and conservation analysis

Sequence alignment of 209 species were manually curated for RasGAP and alignments were automatically generated for RASA2, RASA3, RASA4, RASAL1, RASAL2, RASAL3, SynGAP, and DAB2ip using the Consurf server^58^ by inputting the primary amino acid sequence obtained from UniProt^59^. Conservation was then mapped onto either the AlphaFold prediction or the experimentally determined crystal structure. Sequences aligned using Clustal Omega^60^ and presented in Jalview (v2.11.3.2)^61^.

### Single turnover phosphate sensor GAP assays

Single turnover RasGAP kinetics assays were performed using the Phosphate Sensor system^62,63^ to detect inorganic phosphate (Invitrogen; ThermoFisher Scientific catalogue number PV4407). Assays were performed in 384-well black microplates using a Synergy H1 Microplate Reader with its associated Gen5 software (v3.11.19) (BioTek) in fluorescence mode with an excitation wavelength of 430 nm, an emission wavelength of 450 nm, and bandwidths of 10 nm. Each 20 µL reaction contained 1 µM Phosphate Sensor, 25 nM RasGAP construct, and 0.8 µM HRas•GTP, in 50 mM Tris pH 8, 50 mM NaCl, 10 mM EDTA, 0.01% Triton X-100, 1 mM Tris(2 carboxyethyl) phosphine. Reactions were conducted for 1 hour at 30°C with readings every minute. Data were exported to Excel (v16.66.1) (Microsoft) and normalized according to ^63^. Three or more independent replicates were performed for each condition, each consisting of two duplicates. Data were converted from relative fluorescence units (RFU) to phosphate turnover (µM) via normalization to highest fluorescence signal as per ^35^. Linear fits were applied to the initial linear portion of each curve and slopes were plotted and compared. Statistical significance was determined via ordinary one-way ANOVA with Tukey’s multiple comparisons test.

### Michaelis-Menten single turnover phosphate sensor kinetics assays

Michaelis-Menten single turnover kinetics assays derived from the protocol of ^64^ were performed using the Phosphate Sensor system to detect inorganic phosphate (Invitrogen; ThermoFisher Scientific catalogue number PV4407). Assays were conducted in 384-well black microplates using a Synergy H1 Microplate Reader with Gen5 software v3.11.19 (BioTek) in fluorescence mode with an excitation wavelength of 430 nm, an emission wavelength of 450 nm, and bandwidths of 10 nm. Each 20 µL reaction contained 10 µM Phosphate Sensor, 12.5 mM NaCl, 20 mM Tris pH 8, 5 mM MgCl_2_, 2.5 mM EDTA, 1 mM TCEP, and 0.01% Triton X-100. RasGAP construct concentrations ranged from 5-25 nM. HRas was loaded with GTP and added at concentrations ranging from 10-150 µM, with GTP loading monitored by anion exchange chromatography following protein denaturation^65^. Briefly, HRas was incubated with 100-fold molar excess GTP for ten minutes at 37°C in the presence of 10mM EDTA. Next, 15 mM MgCl_2_ was added, the sample centrifuged at 16,800 rcf for 10 minutes at 4°C, and excess nucleotide removed via size exclusion chromatography on a Superdex 75 Increase 10/300 GL (GE Healthcare) in a buffer containing 20 mM Tris pH 8 and 10 mM EDTA. Fractions containing HRas•GTP were concentrated, frozen in liquid nitrogen, and stored at -80°C until use. To test loading efficiency, 10 nmol HRas was boiled for 15 minutes to promote nucleotide release, centrifuged at 16,800 rcf for 10 minutes at 4°C to remove denatured protein, and the supernatant was applied to a MonoQ column (GE Healthcare) to perform anion exchange chromatography using a continuous NaCl gradient from 0-40% of 1 M NaCl in 20 mM Tris pH 8.5. Peaks corresponding to GDP and GTP were observed via correlation to standards run separately. Peak area was integrated, and loading efficiency was determined by the proportion of GTP to (GDP + GTP). Efficiency range is 75-85%. This efficiency correction was then applied to measured HRas concentrations to account only for HRas•GTP.

Reactions were conducted for twenty minutes at 30°C upon addition of HRas•GTP with readings every twelve seconds. Reactions containing HRas•GTP alone were conducted in parallel to assess the contribution of intrinsic GTPase activity. Five independent replicates were performed for each condition, each consisting of two wells in duplicate which were analyzed together to obtain one Michaelis-Menten fit per replicate. Data were exported to Excel (v16.66.1) (Microsoft) and processed in Prism (v10.1.1) (GraphPad Software, MA). Data were converted from relative fluorescence units (RFU) to phosphate turnover (µM) via a single exponential standard curve produced with varying concentrations of Phosphate Standard (Millipore Sigma 20-103). From the resulting fit, phosphate concentrations could be back calculated from observed RFU values. Linear fits were applied to the initial linear portion of each reaction curve as determined visually. Slopes for curves containing RasGAP were baseline-corrected with slopes for curves containing corresponding concentrations of HRas•GTP only to account only for GAP-mediated GTP hydrolysis. Corrected rates were plotted against HRas•GTP concentration and fit to the Michaelis-Menten equation to obtain k_cat_ and K_M_ with their respective standard deviations. Catalytic efficiency (k_cat_/K_M_) was determined directly from these data. Statistical significance was determined via Brown-Forsythe ANOVA test with Dunnett’s T3 multiple comparisons test.

### Multiple turnover malachite green GAP assays

Multiple turnover endpoint SynGAP assays were conducted using Malachite Green (Millipore Sigma catalogue number MAK307) according to manufacturer protocols in clear, flat-bottomed 96-well microplates. Each 80 µL reaction contained 200 µM GTP, 5 µM SynGAP construct (GAP WT, C2-GAP WT, C2-GAP R401S, or buffer only), 5 µM Rap1b WT, 125 mM NaCl, 60 mM Tris pH 7.5, 5 mM MgCl_2_, 20 mM EDTA, and 0.01% Triton X-100. Five independent replicates were performed for each condition, each consisting of two wells in duplicate which were then analyzed together. Assays were initiated at room temperature by addition of GTP and quenched simultaneously with 20 µL Malachite Green Working Reagent to obtain timepoints at 30 seconds and 10, 20, 30, 40, 50, and 60 minutes. After developing for 30 minutes at room temperature, absorbance at 620 nm (A_620_) was read using a Synergy H1 Microplate Reader with its associated Gen5 software (v3.11.19) (BioTek) in absorbance mode. Data were exported to Excel (v16.66.1) (Microsoft) and processed in Prism (v10.1.1) (GraphPad Software, MA). Data were converted from A_620_ to phosphate turnover (µM) via a linear standard curve produced according to manufacturer protocols using Phosphate Standard (Millipore Sigma). Data were plotted as phosphate turnover vs. time for each construct. Linear fits were applied to each curve and slopes were plotted and compared. Statistical significance was determined via ordinary one-way ANOVA with Dunnett’s multiple comparisons test.

### Generation of Rasa1 R698C mice

CRISPR/Cas9 technology was used to change codon R698 to R698C in the mouse Rasa1 gene. Codon R698 is located in exon 16 of Ensembl.org gene model Rasa1-201 (ENSMUST00000109552.3). The CRISPOR algorithm^66^ was used to identify specific single guide RNAs (sgRNA). sgRNA predicted to cut the chromosome in intron 15 and intron 16 were tested to determine if they cause chromosome breaks. sgRNAs were chemically synthesized with phosphorothioate modifications by Integrated DNA Technologies (IDT)^67,68^. High fidelity Streptomyces pyogenes Cas9 endonuclease protein (HiFi, ^69^) was obtained from (IDT). sgRNAs (60 ng/µl) was complexed with HiFi Cas9 (50 ng/µl) and individually tested to determine if the ribonucleoprotein (RNP) complexes cause chromosome breaks in mouse zygotes. RNPs were microinjected into fertilized mouse eggs. Eggs were placed in culture until they developed into blastocysts. DNA was extracted from individual blastocysts for analysis. PCR with primers spanning the predicted cut site was used to generate amplicons for Sanger sequencing. The process is essentially as described^70^. sgRNA candidates were tested with Forward primer: 5’ CTCCCTTGATACCTCACTAGCTATCTCAA 3’; Reverse primer TCTCTCTGTCATTTAGTGTGCATAACAACA 3’; 955 bp amplicon. Sequence chromatograms of amplicons from individual blastocysts were evaluated to determine if small insertions/deletions caused by non-homologous end-joining (NHEJ) repair of chromosome breaks were present. sgRNA C345A was found to induce chromosome breaks in intron 15. It targets the sequence 5’ TTAAATGCTGTCCACTAAGC (PAM=AGG) 3’ (CFD score of 91^71^). sgRNA C345X to was found to induce chromosome breaks in intron 16. It targets the sequence 5’ TCTTCTCTGCCATTAATATA (PAM=GGG) 3’ (CFD score of 86).

After determining that C345A and C345X cause chromosome breaks, RNPs were combined with 5 ng/µl single stranded Megamer DNA donor (IDT). The sequence of the single stranded oligonucleotide DNA donor was: CACAATCCCATGTTCCCATACTACTACAAAATGCCACATAACAGGCCTTCAGTAAGTGAAGAAAAAAAAGTGAAGTGTTGACTTTGTGGTAGTGTTGAAAAaCTGCTTAGTGGACAGCATTTAAAACTAGAGAAAGGGATTATCAAGAAAAGTTTATTATGTGACTGTAGTGGTTTTCTAATATCGTAATTTGTTTTTTTCTTTGTAG<u>TATTTATGCGTTGCCAGTTGAGCAGGTTACAGAAAGGACATGCAACAGATGAATGGTTTCTACTAAGTTCCCATATACCACTAAAAGGTATTGAACCAGGATCTTTG**tGT**GTTCGAGCACGATACTCCATGGAAAAAATCATGCCAGAAGAAGAGTACAGTGAATTTAAAGAG</u>GTATTGTTTACTAGTTTAATCGTTTTTAATGATACTAAACAGAAATATTTACTATTTAATAAAACTCCATTAAGTTAAATAGTAACAaCCTATATTAATGGCAGAGAAGAAGCCAGATGTAAAAAACACATACTGTGATTTTGATCTTAATGAGACTAGGAATATAAAGTCAATTTTTAACAATTGTATTCgcggccgc (exon 1 is underlined, codon 698 is shown in bold, lower case letters indicate coding changes to the wild type introns to disrupt the PAM needed by Cas9 to make chromosome breaks, a terminal NotI restriction enzyme site was added to improve ssDNA synthesis efficiency). The CRISPR reagents were microinjected into fertilized mouse eggs produced by mating superovulated B6SJLF1 female mice (Jackson Laboratory stock no. 100012) to B6SJLF1 male mice as described^69^. CRISPR/Cas9 microinjection of B6SJLF1 zygotes produced 57 potential founder mice. Six generation zero founder (G0) pups were identified by Sanger sequencing of amplicons spanning exon 16. Forward primer: 5’ ATGCCACATAACAGGCCTTCAG 3’; Reverse primer GCTGAAAGTACACAACAAGACCA 3’; 585 bp amplicon. Two independent G0 founders (mice #s 497 and 554) were mated with *Rasa1* floxed mice on a mixed 129S/v C57BL/6J background^72^ to obtain germline transmission of the *Rasa1* R698C mutant. Targeted amplicon sequencing was used to test all of the top off-target candidates predicted by CRISPOR. No off-target mutations were present. This is in agreement with the findings of Anderson *et al*. ^73^ that the use of high specificity sgRNA and ESPCAS9 makes off-target hit in mouse models highly unlikely. All experiments performed with mice were in compliance with University of Michigan guidelines and were approved by the university committee on the use and care of animals.

### Embryonic development of Rasa1 R698C mice

Analysis of the effect of the R698C mutation upon embryonic development was assessed for both independent line. Genotypes of P6 pups and embryos at different stages of development from crosses of heterozygous parents were determined by PCR of tail genomic DNA using the same exon 16 flanking primers as above, followed by Sanger sequencing. Whole embryos were imaged using a Leica M80 dissecting microscope fitted with a DMC4500 camera (Leica). Whole E9.5 embryo lysates were prepared in RIPA buffer, and amounts of RASA1 and actin in lysates were determined by Western blotting as described^72^.

### Vascular development in Rasa1 R698C mice

E9.5 embryos and E10.5 yolk sacs were fixed in 1% paraformaldehyde for 1 hour and embryos were cleared using the PEGASOS method as described (ref). Embryos and yolk sacs were blocked in 3% donkey serum/0.3% Triton X-100 in PBS overnight and stained with anti-CD31 antibody (MEC13.3, eBioscience) in 0.3% Triton X-100 in PBS overnight. Embryos were then incubated with Alexa Flour 488 donkey anti-rat Ig overnight before mounting and viewing on a Leica SP5 X confocal microscope (Leica Microsystems).

### Phospho-MAPK staining

Fixed E9.5 embryos were embedded in paraffin and 5 µm sections were stained with antibodies against CD31 (SZ31, Dianova) and phospho-ERK MAPK (D13.14.4E, Cell Signaling Technology) and Hoechst (Invitrogen) as described (PMID: 35015735) Sections were viewed on a Bx60 upright fluorescence microscope (Nikon).

### Generation of Rasa1 T603A mice

The same CRISPR/Cas9 methods as above were used to change codon T603 to T603A in exon 14 the mouse Rasa1 gene. sgRNAs predicted to cut the chromosome in intron 13 and intron 14 were tested to determine if they cause chromosome breaks. sgRNA candidates were tested with C344A Forward primer: 5’ AATCAGAGCTAATATCGTGAATCCCTTGG 3’; Reverse primer CTTACTCAAAGACAAACTCCTCTGACCAT 3’; 671 bp amplicon and C344Z Forward primer: 5’ TGGTGTCTTAGAATACCTTAACCATGAGAC 3’; Reverse primer TAGCATGTGTTTCACTTCCATTTTGACAC 3’; 866 bp amplicon. sgRNA C344A was found to induce chromosome breaks in intron 13. It targets the sequence 5’ TGTAGTTTTCATTTGGATGA (PAM=TGG) 3’ (CFD score of 75, Doench) in the reverse direction. sgRNA C344Z to was found to induce chromosome breaks in intron 14. It targets the sequence 5’ ACAGATTATATATAAACAGT (PAM=AGG) 3’ (CFD score of 88) in the reverse direction.

After determining that C344A and C344Z cause chromosome breaks, RNPs were combined with 5 ng/µl single stranded Megamer DNA donor (IDT) and microinjected into fertilized mouse eggs as above. The sequence of the DNA donor was: CTAGAATAGTGAAATCTTAATGTAAGCATATGGTTGCTTATACTGTACATTACATGTAAGCATGTTTATTCCATGTTTGTAGTTTTCATTTGGATGATGtTAGTTGATTAGCAAAGCTTATGTTTTCTTTTAAATATTAAGGAAAAAAGTCTTTTTTTTTTTTATTTAAAAGCCTCATATTCAGGTCTTGAAATGTTGGATATGAATTTTTGTGTTCTCTTTAG<u>GTCAGCAGCCTTGTTTTACATATTGAAGAAGCCCATAAACTACCAGTAAAACACTTT**gCT**AATCCATATTGTAACATCTACTTGAATAGCGTCCAAGTAGCAAAAACTCATGCAAGGGAGGGGCAAAACCCAGTATGGTCAGAGGAGTTTG</u>TCTTTGAGTAAGTCATATCTTGTCATATTCAATCAGTTTTTACAATCATAATTACATTTTCCTaCTACTGTTTATATATAATCTGTTTAAGTTTCTATTGTTTTCTAGGTGTTTTAATAGTATTTTCTCTTAGCTTCTTTCATATTTGAATTACTC. (exon 14 is underlined, codon 603 is shown in bold, lower case letters indicate coding changes to the wild type introns to disrupt the PAM needed by Cas9 to make chromosome breaks, a terminal NotI restriction enzyme site was added to improve ssDNA synthesis efficiency).

CRISPR/Cas9 microinjection of zygotes produced 41 potential founder mice. One generation zero founder (G0) pup was identified by Sanger sequencing of amplicons spanning exon 14. Forward primer: 5’ GGAGAGGCTCCGAGACTGACTTGT 3’; Reverse primer CAGGAGGAAGATCACTAAAGAAGCA 3’; 875 bp amplicon. The founder was mated with *Rasa1* floxed mice on a mixed 129S/v C57BL/6J background^72^ to obtain germline transmission of the *Rasa1* T603A mutant. Genotypes of P6 pups from crosses of heterozygous parents were determined by PCR of tail genomic DNA using the same exon 14 flanking primers as above, followed by Sanger sequencing.

## Data availability

Atomic coordinates and structure factors for the refined structural models described in this paper have been deposited in the Protein Data Bank (PDB) under accession code 9BZ4. X-ray diffraction images are available online at SBGrid Data Bank: doi:10.15785/SBGRID/1107). Source data are provided with this paper.

## Research animal ethical compliance statement

All experiments performed with mice complied with University of Michigan guidelines and were approved by the university committee on the use and care of animals.

## Acknowledgements

We thank Nalini Natarajan, Karen Anderson, Enrique De La Cruz, and Moitrayee Bhattacharyya for helpful discussions. We thank Jim Murphy for training and support. We thank the beamline scientists of NE-CAT at the Advance Photon Source. We acknowledge Wanda Filipiak and Galina Gavrilina and the Transgenic Animal Model Core of the University of Michigan’s Biomedical Research Core Facilities for assistance with the design and production of RASA1 R698C and T603A knockin mice. This work is based upon research conducted at the Northeastern Collaborative Access Team beamlines, which are funded by the National Institute of General Medical Sciences from the National Institutes of Health (P30 GM124165). The Eiger 16M detector on the 24-ID-E beam line is funded by a NIH-ORIP HEI grant (S10OD021527). This research used resources of the Advanced Photon Source, a U.S. Department of Energy (DOE) Office of Science User Facility operated for the DOE Office of Science by Argonne National Laboratory under Contract No. DE-AC02-06CH11357. M.E.P. supported by T32GM007324 and F31HL167578. E.D.H., Z.T.F., and T.L.S. supported by P30CA046592. K.J.V. supported by T32GM008283 and F31HL165968. This research was supported by NIH grants HL120888 and HL146352 to P.D.K. and NS117609, GM102262 and GM138411 to T.J.B.

## Author contributions

Conceptualization, Methodology, Writing: M.E.P., D.C., P.D.K., and T.J.B. Investigation, Data Curation, Visualization: M.E.P., D.C., K.J.V., E.H., Z.T.F., N.L.L., T.L.S., and A.L.S. Supervision:

P.D.K. and T.J.B.

Competing interests

The authors declare no competing interests.

## Materials and Correspondence

P.D.K. and T.J.B.

