## Supplemental Figures S1-S10 and Tables S1-S6 for "The C2 domain augments Ras GTPase Activating Protein catalytic activity"

**Table S1. Single turnover RasGAP assay.** Single turnover GAP assay. Initial rates for malachite green reporting on phosphate release from GTP loaded Ras. RasGAP constructs: C2-GAP and GAP alone. Related for Figure 1b. SD, standard deviation.

| **RasGAP Construct** | **Initial Rate ($min^{-1}$)** | **Initial Rate SD ($min^{-1}$)** |
|---|---:|---:|
| C2-GAP | 0.03565 | 0.003782 |
| GAP | 0.02383 | 0.003438 |
| None (HRas only) | 0.01643 | 0.004375 |

**Table S2. Crystallographic statistics.** Statistics for the highest resolution shell are shown in parentheses. RMSD: root-mean-square deviation.

| **Data Collection** | **RasGAP C2-GAP (residues 588-1044)** |
|---|---|
| PDB Accession Code | 9BZ4 |
| Wavelength (Å) | 0.97918 |
| Resolution range (Å) | 87.5 - 2.45 (2.5 - 2.45) |
| Space group | *P* 1 |
| Unit cell dimensions (Å) | a=70.9, b=94.0, c=96.7 |
| α, β, γ (°) | α=95.8, β=111.1, γ=108.5 |
| Total observations | 1,157,715 |
| Unique reflections | 75,708 (7,149) |
| Multiplicity | 10.0 (7.0) |
| Completeness (%) | 96.1 (90.5) |
| Mean $I/\sigma(I)$ | 12.1 (1.7) |
| Wilson B-factor (Å$^2$) | 61.4 |
| $R_{merge}$ (%) | 30.8 (149.5) |
| $R_{meas}$ (%) | 34.0 (172.8) |
| $R_{pim}$ (%) | 10.8 (64.1) |
| CC½ | 98.9 (47.6) |
| **Refinement** | |
| Resolution range (Å) | 87.5-2.45 (2.47-2.45) |
| Reflections used in refinement | 75,687 (2,102) |
| Reflections used for $R_{free}$ | 3,879 (122) |
| $R_{work}$ (%) | 22.8 (35.0) |
| $R_{free}$ (%) | 28.4 (43.0) |
| Number of non-hydrogen atoms | 14,329 |
| Macromolecules | 14,303 |
| Ligands | 0 |
| Solvent | 26 |
| Protein residues | 1,774 |
| **Residue numbers built** | |
| Chain A | 588-781, 787-978, 984-1044 |
| Chain B | 588-782, 790-835, 838-983, 986-1044 |
| Chain C | 588-663, 666-826, 838-916, 921-1043 |
| Chain D | 588-661, 665-781, 784-836, 842-946, 951-1043 |
| RMSD (bonds, Å) | 0.003 |
| RMSD (angles, °) | 0.57 |
| Ramachandran favored (%) | 97.9 |
| Ramachandran allowed (%) | 2.1 |
| Ramachandran outliers (%) | 0.0 |
| Rotamer outliers (%) | 0.73 |
| MolProbity clashscore | 7.57 (98th percentile) |
| *B*-factors: Average (Å$^2$) | 77.6 |
| Macromolecules (Å$^2$) | 77.7 |
| Chains: A, B, C, D (Å$^2$) | 72.8, 75.9, 79.1, 83.1 |
| Solvent (Å$^2$) | 60.5 |
| Number of TLS groups | 14 |

**Table S3. Michaelis-Menten parameters from single turnover phosphate release assay for RasGAP C2-GAP.** Mutations indicated. SD, standard deviation.

| **RasGAP Construct** | **$k_{cat}$ Mean ($s^{-1}$)** | **$k_{cat}$ SD ($s^{-1}$)** | **$K_M$ Mean (μM)** | **$K_M$ SD (μM)** | **Catalytic Efficiency Mean ($M^{-1}s^{-1}$)** | **Catalytic Efficiency SD ($M^{-1}s^{-1}$)** |
|---|---|---|---|---|---|---|
| GAP | 2.902 | 0.5898 | 80.4 | 21.43 | 37164 | 7527 |
| C2-GAP | 15.07 | 2.655 | 33.14 | 2.814 | 457580 | 91197 |
| C2-GAP R707S | 0.7917 | 0.3666 | 89.78 | 56.51 | 9865 | 2990 |
| C2-GAP R707D | 0.7852 | 0.3318 | 183 | 116.2 | 4703 | 1531 |

**Table S4. Michaelis-Menten parameters from single turnover phosphate release assay for RasGAP ΔN.** Mutations indicated. SD, standard deviation.

| RasGAP Construct | $k_{cat}$ Mean ($s^{-1}$) | $k_{cat}$ SD ($s^{-1}$) | $K_M$ Mean (μM) | $K_M$ SD (μM) | Catalytic Efficiency Mean ($M^{-1}s^{-1}$) | Catalytic Efficiency SD ($M^{-1}s^{-1}$) |
|---|---|---|---|---|---|---|
| ΔN WT | 8.686 | 1.875 | 73.32 | 18.56 | 121322 | 25177 |
| ΔN R707S | 2.076 | 1.23 | 295.7 | 178.6 | 7076 | 662.6 |
| ΔN R707C | 1.08 | 0.5135 | 122.1 | 78.72 | 9528 | 2642 |
| ΔN W689F | 2.122 | 0.4236 | 77.88 | 26.76 | 28968 | 8045 |

**Table S5. SynGAP enzyme assay data.** SD, standard deviation.

| SynGAP Construct | Initial Rate (µM/min) | Initial Rate SD (µM/min) |
|---|---|---|
| GAP | 0.001811 | 0.009611 |
| C2-GAP | 0.1874 | 0.008362 |
| C2-GAP R401S | -0.02304 | 0.01538 |
| None (Rap1b only) | -0.004808 | 0.01027 |

**Table S6. Cancer associated missense mutations of conserved residues in the C2 domain of RasGAP family proteins.** Analysis of mutations conducted using cBioPortal [74-76] and COSMIC [77].

| Mutation | Cancer type | Citation |
|---|---|---|
| **RASA1** | | |
| H604N | Uterine | Nguyen *et al.* [78] |
| W689R | Lung | Curated Study, ICGC(LUSC-KR), study ID: COSU583 [77,79] |
| S705F | Glioma | Lee *et al.* [80] |
| S705P | Pancreatic | TCGA research network – PAAD pan-cancer atlas 2018 |
| R707C | Colon | Nguyen *et al.* [78] |
| R707C | Colorectal | Nguyen *et al.*, [78] |
| R707C | Ovarian | Nguyen *et al.* [78] |
| R707H | Colorectal | Giannakis *et al.* [81] |
| R707H | Stomach | Nguyen *et al.*, [78] |
| R707H | Urothelial | Nguyen *et al.* [78] |
| R707P | Uterine | TCGA research network – UCEC pan-cancer atlas 2018 |
| **RASA2** | | |
| R310L | Lung | TCGA research network – LUAD pan-cancer atlas 2018 |
| R310Q | Colorectal | Giannakis *et al.* [81] |
| R310Q | Lung | TCGA research network – LUSC pan-cancer atlas 2018 |
| R310Q | Squamous skin | Chang *et al.* [82] |
| R310Q | Uterine | TCGA research network – UCEC pan-cancer atlas 2018 |
| **RASA3** | | |
| S282F | Squamous skin | Pickering *et al.*[83] |
| S282F | Melanoma | Liu *et al.* [84] |
| R284Q | Esophageal | Zhang *et al.* [85]; Cheng *et al.* [86]; Curated Study, ICGC(ESCA-CN), study ID: COSU582 [77,79] |
| R284W | Colorectal | Giannakis *et al.*[81] |
| **RASAL1** | | |
| A249V | Glioblastoma | Barthel *et al.* [87] |
| A249V | Head and neck squamous cell | TCGA research network – HNSC pan-cancer atlas 2018 |
| R251Q | Lung | TCGA research network – LUSC pan-cancer atlas 2018 |
| **SynGAP** | | |
| R272W | Squamous skin | Chang *et al.* [82] |
| R401Q | Melanoma | Snyder *et al.* [88]; Miao *et al.* [89] |
| R401W | Oligodendroglioma | ICGC/TCGA Consortium[90]<br>TCGA research network – LGG pan-cancer atlas 2018 |
| **RASAL2** | | |
| S283Y | Cutaneous melanoma | Van Allen *et al.*[91] |
| S283Y | Lentigo maligna melanoma | Miao *et al.* [89] |
| R285Q | Uterine | TCGA research network – UCEC pan-cancer atlas 2018 |
| **RASAL3** | | |
| W403C | Stomach | Curated Study, ICGC(GACA-JP), study ID: COSU683 [77,79] |
| R416Q | Melanoma | Liu *et al.* [84] |

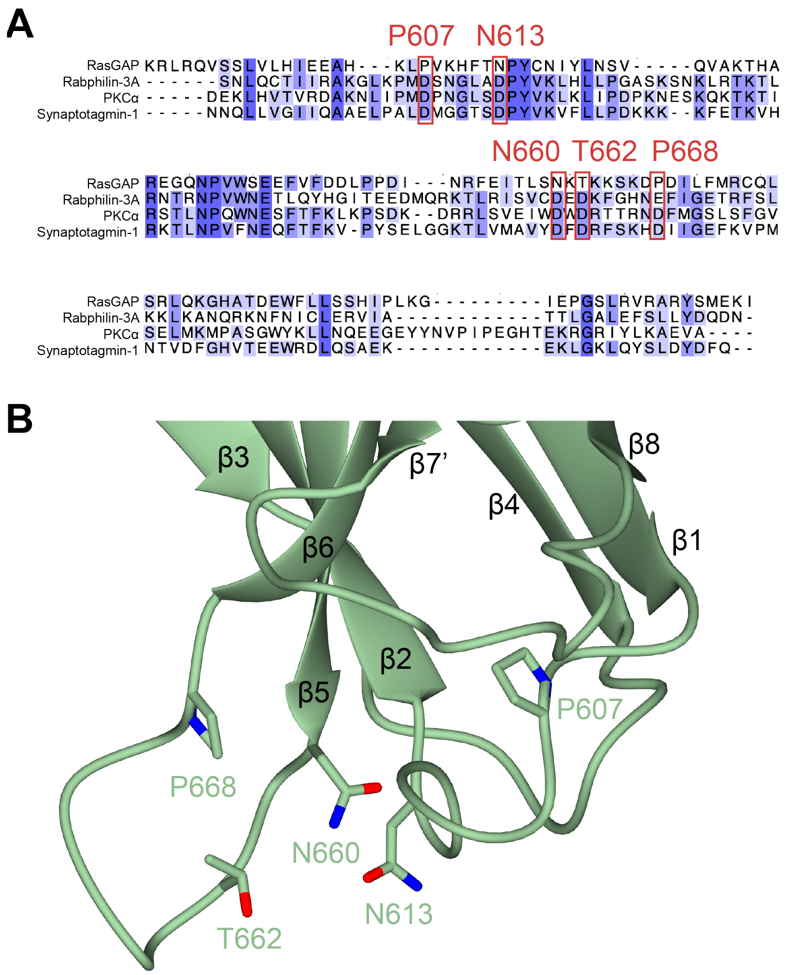

**Figure S1. RasGAP C2 domain does not contain a calcium binding site. A)** Alignment of RasGAP C2 calcium binding residues and the canonical C2 domains Rabphilin-3A, PKCα, and Synaptotagmin-1 [37]. Colors represent percent identity, with darker blue indicating greater conservation. Canonical calcium binding residues are conserved acidic residues at the indicated positions. RasGAP residues at these positions are indicated. **B)** Calcium binding-equivalent residues of the structure of RasGAP. These residues are uncharged and distant in space, suggesting a lack of calcium binding ability. Secondary structure features are indicated.

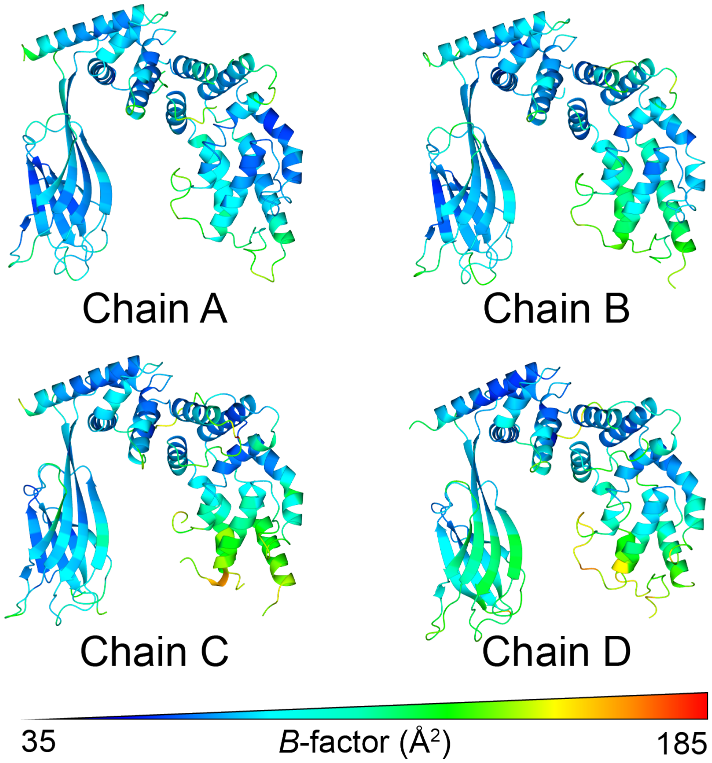

**Figure S2. Four copies of RasGAP C2-GAP in the asymmetric unit.** Four copies of RasGAP C2-GAP in the crystal asymmetric unit, aligned to the GAP domain catalytic core. Colored by *B*-factor from low (blue) to high (red).

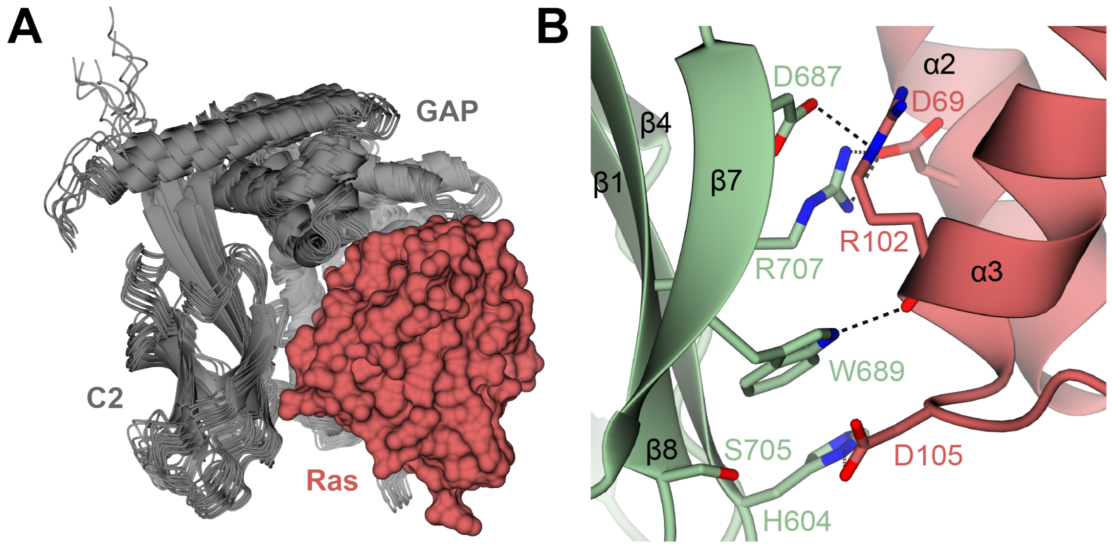

**Figure S3. AlphaFold models of the complex of RasGAP C2-GAP with Ras. A)** Overlay of seventeen AlphaFold multimer models [41] of the C2-GAP region of RasGAP (gray ribbons) with Ras (red; representative surface shown) aligned to Ras $C_\alpha$ atoms. All models show similar domain arrangement and predict direct contact between C2 and Ras. **B)** Close-up of representative AlphaFold multimer model. Highly conserved residues on the surface of the C2 domain of RasGAP (green) and their contacts with Ras (red) are shown. Polar contacts are shown as black dashed lines. Secondary structure features are indicated.

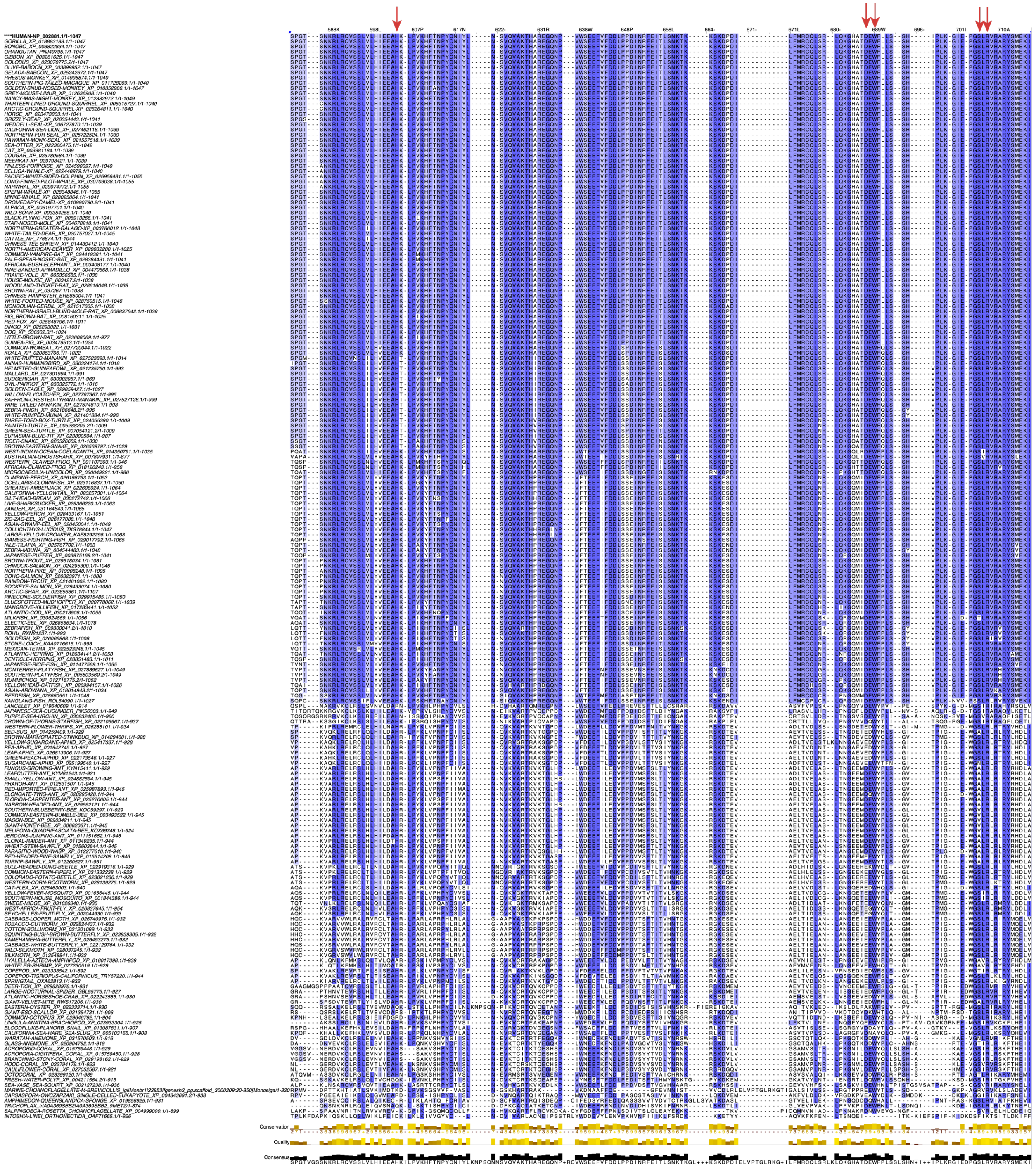

**Figure S4. Sequence alignment of RasGAP.** Alignment of residues 582-717 of RasGAP, approximately comprising the C2 domain. Colors represent percent identity, with darker blue indicating greater conservation. Conserved residues on the putative Ras-binding face of the C2 domain are indicated with red arrows. This contiguous patch comprises residue H604 of strand β1, D687 and W689 of strand β7, and S705 and R707 of strand β8.

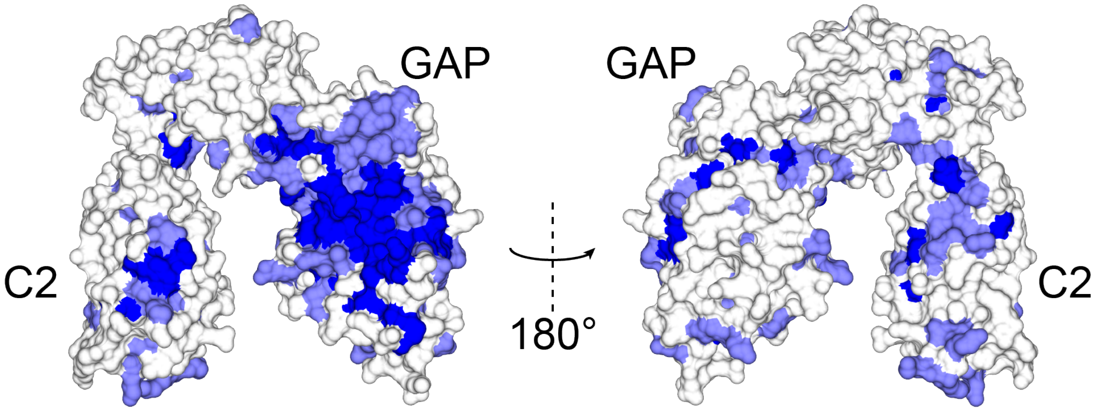

**Figure S5. Conservation of the C2-GAP region of RasGAP.** Alignment of 209 sequences of RasGAP from humans to sponges (**Figure S4**) with conservation mapped onto the structure using the Consurf server [58]. Blue indicates complete conservation and white low conservation. GAP and C2 domains indicated.

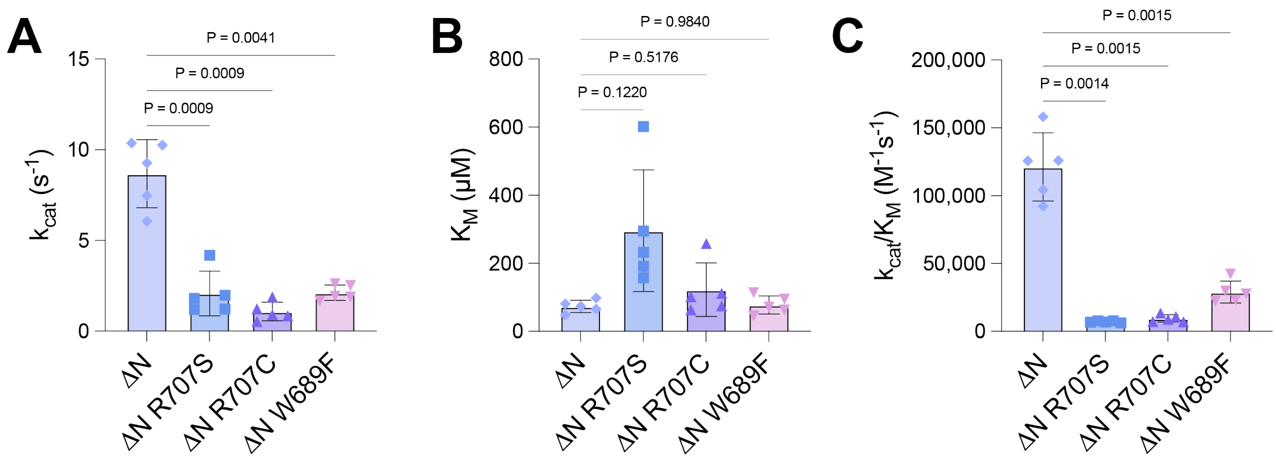

**Figure S6. Michaelis-Menten parameters from single turnover phosphate release assay for RasGAP ΔN. A)** $k_{cat}$ of RasGAP ΔN constructs. Bars indicate mean ± SD with replicates (n=5) shown. *P* values for relevant comparisons are shown. Statistical significance determined via Brown-Forsythe ANOVA test (*F* = 44.70, 11.120 degrees of freedom) with Dunnett's T3 multiple comparisons test. **B)** $K_M$ of RasGAP constructs. Bars indicate mean ± SD with replicates (n=5) shown. *P* values for relevant comparisons are shown. Statistical significance determined via Brown-Forsythe ANOVA test (*F* = 5.593, 8.806 degrees of freedom) with Dunnett's T3 multiple comparisons test. **C)** Catalytic efficiency ($k_{cat}/K_M$) of RasGAP constructs. Bars indicate mean ± SD with replicates (n=5) shown. *P* values for relevant comparisons are shown. Statistical significance determined via Brown-Forsythe ANOVA test (*F* = 82.49, 7.910 degrees of freedom) with Dunnett's T3 multiple comparisons test. RasGAP ΔN construct encompasses all folded domains.

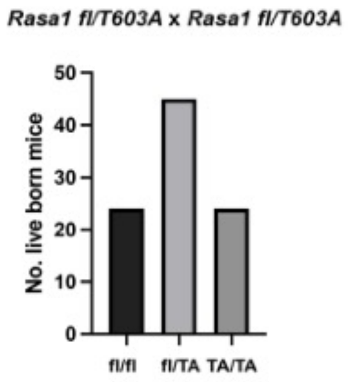

**Figure S7. A *Rasa1 T603A* mutation does not cause embryonic lethality in homozygous form.** *Rasa1 fl/T603A* heterozygous mice were crossed and numbers of live-born pups of the indicated genotypes were determined. Differences are not statistically significant and determined with a Chi-squared test.

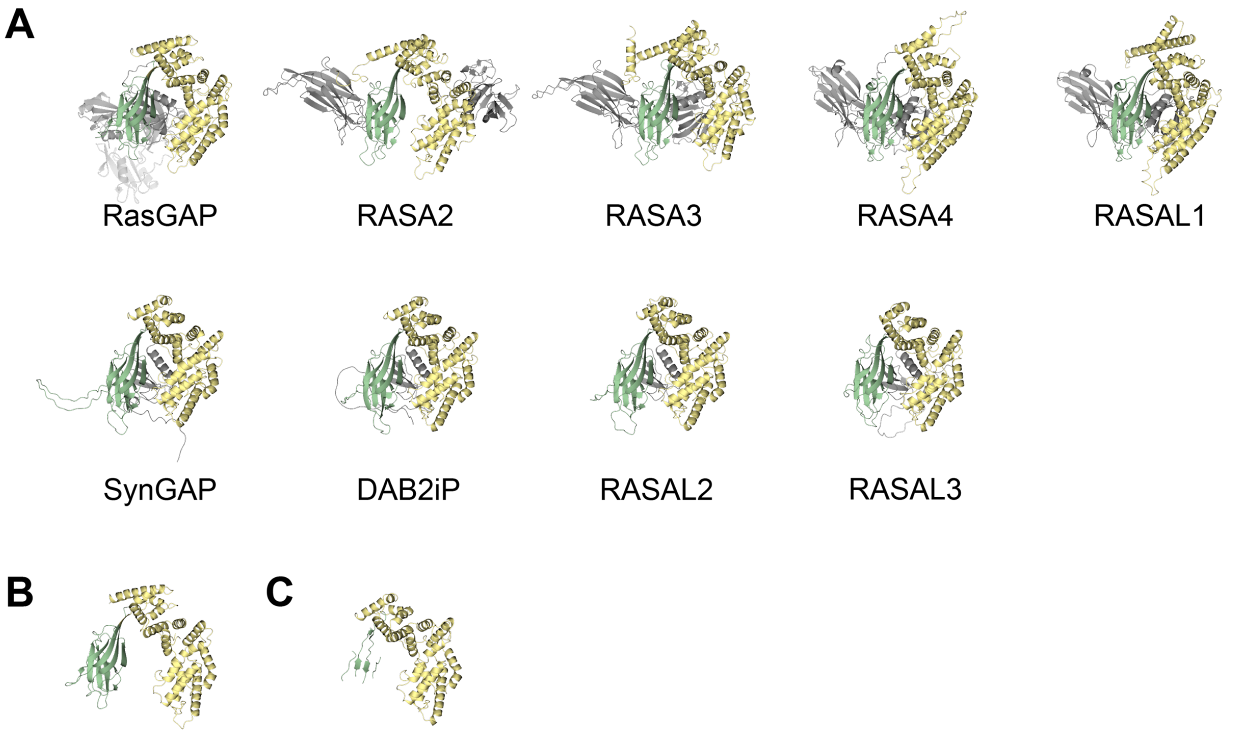

**Figure S8. AlphaFold predictions of nine human GAP proteins for Ras. A)** AlphaFold predictions for the nine human GAP proteins for Ras containing a C2 domain. RASA1, AF-P20936-F1; RASA2, AF-Q15283-F1; RASA3, AF-Q14644-F1; RASA4, AF-O43374-F1; RASAL1, AF-O95294-F1; RASAL2, AF-Q9UJF2-F1; RASAL3, AF-Q86YV0-F1; SynGAP, AF-Q96PV0-F1; DAB2ip, AF-Q5VWQ8-F1. RASA1 is version from 01-JUL-21, all others are from 01-JUN-22. GAP domains colored yellow, proximal C2 domains colored green, other domains colored gray. **B)** Crystal structure of RASA1 C2-GAP (this paper, PDB accession code: 9BZ4). **C)** Crystal structure of SynGAP C2-GAP construct[28] (PDB accession code: 3BXJ).

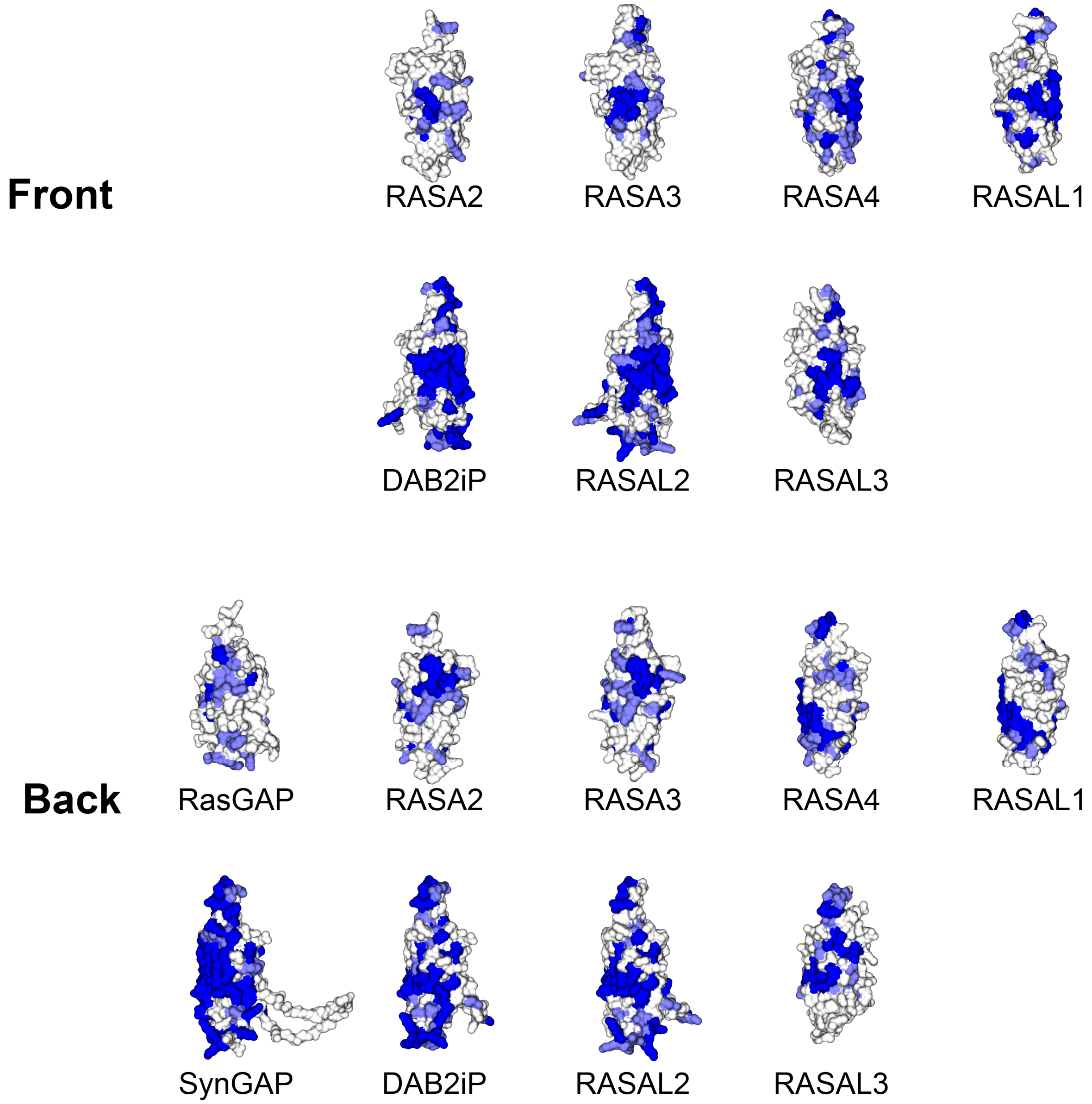

**Figure S9. Sequence conservation mapped onto AlphaFold prediction of GAP-proximal C2 domains.** For each of the nine GAPs for Ras sequence conservation is mapped onto the predicted GAP-proximal C2 domain. Front face of C2 beta sandwich comprises strands β7, β8, β1, and β4. Back face comprises strands β3, β2, β5, and β6. High conservation colored blue, low conservation colored white. RasGAP, crystal structure; RASA2, AF-Q15283-F1; RASA3, AF-Q14644-F1; RASA4, AF-O43374-F1; RASAL1, AF-O95294-F1; RASAL2, AF-Q9UJF2-F1; RASAL3, AF-Q86YV0-F1; SynGAP, AF-Q96PV0-F1; DAB2ip, AF-Q5VWQ8-F1. Models are from AlphaFold version 01-JUN-22. Also see Figure 4.

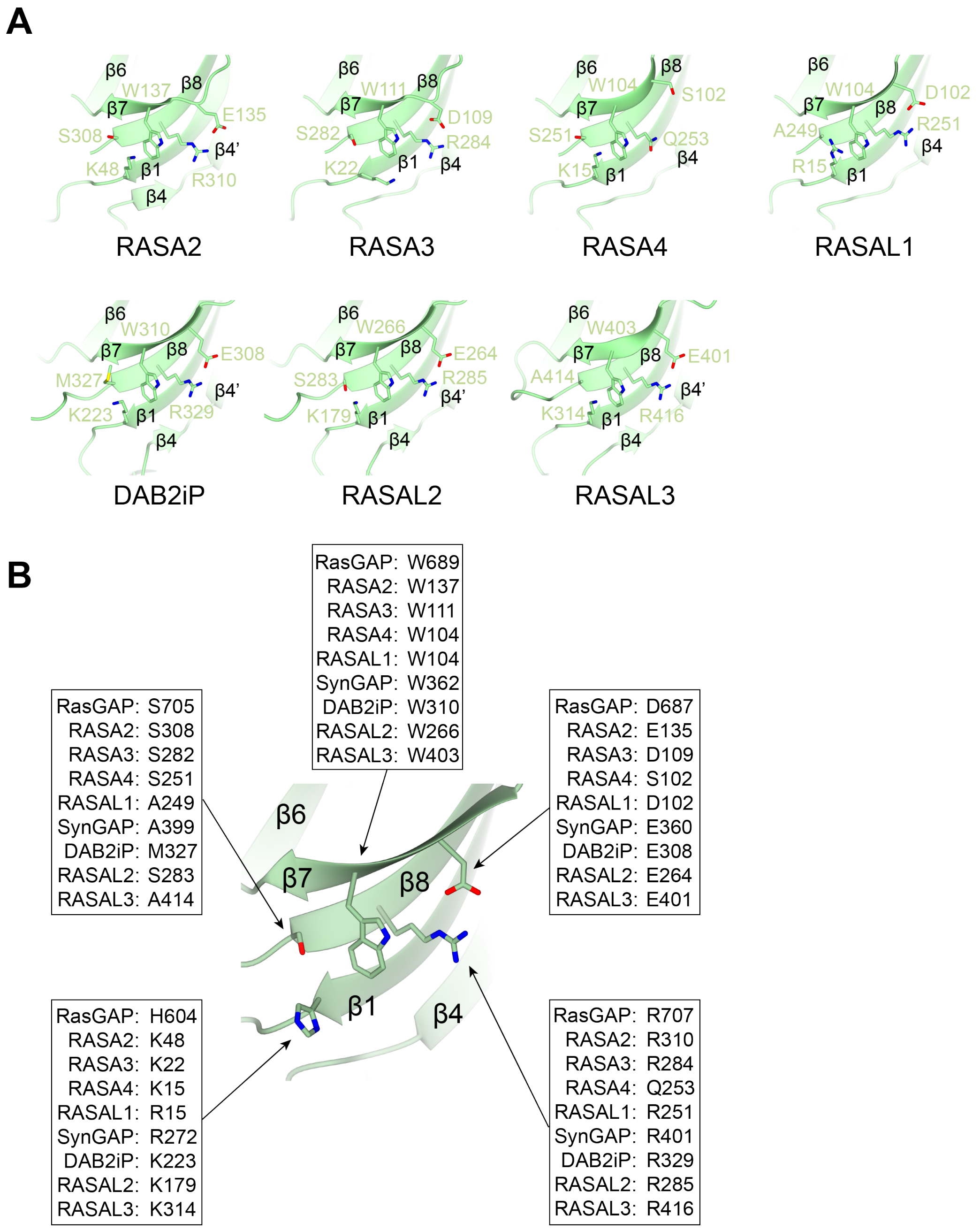

**Figure S10. Closeup of conserved C2 surface in AlphaFold models of GAP proteins for Ras. A)** Closeup of AlphaFold predictions of the C2 domains of GAP proteins for Ras. Evolutionarily conserved C2 domain residues are shown for RASA2, RASA3, RASA4, DAB2iP, RASAL2, and RASAL3. Those for RasGAP, SynGAP and RASAL1 are shown in Fig 4b. Secondary structure features indicated. **B)** Schematic of equivalent conserved residues on C2 surface of RasGAP family members illustrated on the crystal structure of RasGAP. Secondary structure features indicated.
